# Testing Sensorimotor Timing in a Living Laboratory – Behavioral Signatures of a Neural Oscillator

**DOI:** 10.1101/2025.06.25.661347

**Authors:** Hélène Serré, Keith Harrigian, Se-Woong Park, Dagmar Sternad

## Abstract

Rhythmic ability has been studied for more than a century in laboratory settings testing timed finger taps. While robust results emerged, it remains unclear whether these findings reflect behavioral limitations in realistic scenarios. This study tested the synchronization-continuation task in a museum with 455 visitors of a wide variety of ages (5-74yrs), musical experiences (0-40yrs) and educational and cultural backgrounds. Adopting a dynamic system’s perspective, three metronome pacing periods were anchored around each individual’s preferred tempo, and 20% faster and 20% slower. Key laboratory findings were replicated and extended: timing error and variability decreased during childhood and increased in older adults and were lower, even with moderate musical experience. Consistent with an oscillator perspective, timing at non-preferred tempi drifted toward their preferred rate. Overall, these findings demonstrate that timing limitations may reflect attractor properties of a neural oscillator and its signature is still present even in noisy, naturalistic settings.

## Introduction

The human sense of timing is one of the oldest phenomena that garnered interest in experimental psychology dating back to Stevens, Scripture and others in the 19^th^ century^1–3^. Over more than 100 years, a plethora of studies scrutinized how humans time their motor commands, specifically in rhythmic timing, as it underlies the human fascination with music^4^. Legions of experiments have employed the extremely simple movement – finger tapping -to test the ability to react, predict and entrain to auditory or visual stimuli. In 1886, Stevens introduced the synchronization-continuation paradigm, where participants tap their index finger in synchrony with a metronome^3^. A synchronization segment is followed by a segment where the external cue is withheld, and the participant continues tapping at the same pace^5–10^. This finger tapping task has remained a widely-used experimental paradigm until the present day^11,12^, including neuroimaging in humans and animals^13,14^ to address a large variety of questions. Different models have been proposed to account for the findings, from Wing and Kristofferson’s clock-motor timing^15^ to network and oscillator perspectives^16^.

This body of experimental research has accumulated a considerable number of observations, grounded in a relatively confined set of variables. The main measurements are the time difference or error between the beat and its corresponding tap, and the variability of the inter-tap interval (ITI)^15^. This paradigm has revealed many fine-grained features of time production. The present study re-examined these features in a large diverse cohort in a “living laboratory”, testing the robustness of these observations in a real-world setting, an outreach activity in the Museum of Science in Boston. Our study involved a very diverse sample of museum visitors of all ages, musical experience, educational, professional, cultural and ethnic background. Our specific approach was grounded in a dynamic systems perspective, positing that the behavioral rhythm emerges from a neuromechanical oscillator with attractor properties. We therefore adapted the experimental design to individualize the timing protocol to participants’ preferred period, or spontaneous motor tempo (SMT) and define pacing periods in relation to their SMT. To set the stage for our study, the main results from the previous plethora of studies are summarized.

### Spontaneous Motor Tempo

Sensorimotor synchronization is least variable when moving at one’s preferred rhythm, often called spontaneous motor tempo (SMT)^7,9,17^. The SMT naturally exhibits inter-individual differences determined by biomechanical properties of the limbs^7,18,19^, physiological factors, such as energy consumption^20,21^, the tapping force^22^, and the time of day^23,24^. A stable SMT is also observed in musicians, even though they are trained to synchronize with many different tempi^5^. Further, the SMT slows down with age and musical training^9,10,25^. Although many studies have collected participants’ SMT, only a few have normalized the range of periods to the SMT^5,7^. We anchored the metronome intervals around the SMT, thereby relating people’s timing performance to their own preferred period. This approach is motivated by a dynamic systems perspective assuming that the SMT is the natural period of a neural oscillator. From this vantage point, we hypothesize that the preferred tempo is a stable attractor. When oscillating at a different period, but within the oscillator’s basin of attraction, participants perform with more variability (less stability) and drift towards their preferred period.

### Effects of Musical Experience

Many studies were motivated by how musical practice affects the sense of time and tested professionals with many years of extensive training^26,27^. Unsurprisingly, sensorimotor synchronization is enhanced in musicians due to their natural aptitude and musical training^28^. The extensive practice fine-tunes the neural mechanisms underlying timing and coordination, leading to superior motor timing, documented in lower ITI variability and shorter asynchronies, i.e., the tendency to tap prior to the metronome beat^28–32,33^.

Beyond synchronization, maintaining a steady tempo without cues is also considered a significant quality when playing music. Despite its importance for musicians, this drift in the continuation segment has received very little attention in musically-trained individuals^5^. We hypothesize that musical experience may strengthen the attractor properties attenuating the drift towards their preferred period.

### Effects of Age

Not unexpectedly, older adults (>50 years old) are more variable than younger adults^9^. Synchronization abilities improve significantly as children grow up and typically reach adult levels by 8 to 10 years of age^9,10,25,33,34^. Before that age, ITI variability is higher and asynchrony is longer. The SMT also evolves with age, particularly exhibiting a slowing in older adults. Also, children drift toward their SMT faster than adults^9^. However, these drifts were calculated in reference to the same set of intervals across all participants, not scaled to their preferred period. Our study sets the metronome periods with respect to the individually preferred period, and we hypothesize to see differences in the drift across age.

This brief review of findings presents our current understanding of timing. However, it is noteworthy that these effects are very subtle. The ITI variability (coefficient of variation) in 5-75 year-olds varies from 0.2 in the youngest participants to 0.07 in adults, measured for intervals between 200ms and 1600ms^9,35^. The difference in mean asynchrony between professional musicians and non-musicians is maximally 40ms^29^. These results are typically obtained from tightly controlled experiments, often conducted in sound-proof cabins with precisely tuned tapping keys, and with participants that can follow precise instructions and are usually attentive and receive financial reward for their participation. While these procedures have yielded reliable data and revealed interesting features of timing and their limitations, such scenarios are a far cry from realistic environments^32,36^. Hence, the question arises whether these timing features are strong enough to survive or even matter in the complex uncontrolled real-world environments.

The present study draws upon hundreds of hours of data collected in the environment of an outreach activity in the Museum of Science in Boston. Using a large inter-generational pool of museum visitors who volunteered for a finger tapping task with a synchronization-continuation design, the current study pursued three aims: 1) test the robustness of previously reported basic timing effects in a noisy real-world environment, 2) examine the effects of age in a diverse sample of the population, and 3) investigate the effect of average musical experience. An oscillator perspective motivated the experimental design.

Museum visitors completed a finger tapping task with synchronization and continuation segments in three different conditions: at their spontaneous motor tempo (SMT), at a tempo 20% faster and 20% slower than their SMT. The data collection was performed in an open space of the larger museum area, where museum goers passed by and were spontaneously recruited to participate and learn about ongoing research. Challenges in recording the tapping force data were overcome by a novel algorithm that sensitively detected contacts despite a high degree of variation and noise. Results confirmed that ITI error and variability improved during development and deteriorated for older adults, both in synchronization and continuation segments. Results further showed that musical experience consistently improved both sensorimotor synchronization and rhythm maintenance. Our specific hypotheses on drift were also supported: participants indeed drifted towards their preferred period. However, these drifts did not differ between musicians or across age. Despite this lack of differentiation, our results demonstrated that even in the little controlled conditions of a museum environment, the main timing effects stood the test of time. The findings are discussed in the light of previous studies and through the lens of the dynamic systems’ theoretical framework.

## Results

The experiment was part of the Living Laboratory program of the Museum of Science in Boston, an initiative designed to educate the public through active involvement in scientific experiments. All subjects in this experiment were museum visitors and data collection occurred during 4-hour exhibitions set up every weekend over 8 months. The experiment was conducted by a total of 9 experimenters rotating across the 26 exhibit sessions. Each data collection session required the team work of 3 students—one for recruiting museum visitors and obtaining informed consent, one for providing information about ongoing research and administering a short questionnaire about age and muscial training, and one for operating the experimental apparatus and collecting data. Participants were asked to tap their finger to a metronome for 15s and maintaining the pace for 25s after the metronome terminated. The data collection for each subject was limited to 15 minutes per museum policy. A total of 335 volunteers participated in the study (180 female, 155 male, ages 5-68). Given the uncontrolled nature of the experimental data collection (see Figure 1), some data had to be eliminated from further analysis, ending with a total of 266 subjects (149 females, 117 males, ages 6-68).

**Figure 1:**
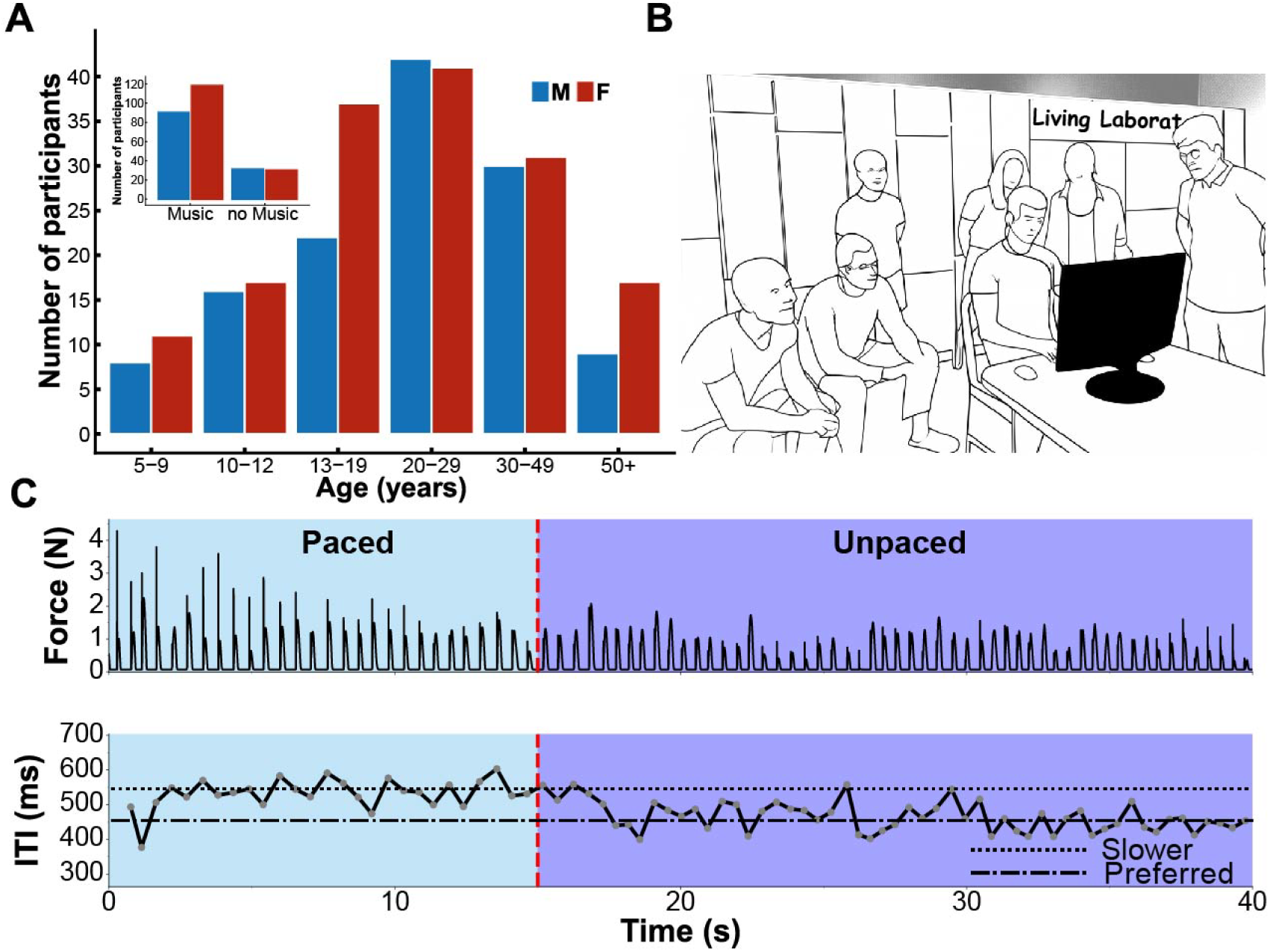
**A**: Distributions of 266 participants of age and gender. The insets shows the split by musical experience. **B**: Experimental data collection at the Museum of Science. **C**: Example of a tapping trial in the slower condition. The two shades separated by the red dashed line highlights the paced and unpaced segments. Top: Raw force signal. Bottom: Inter-tap intervals (ITIs) clustered around the metronome beats (550 ms). A drift toward the preferred period (460 ms, dashed line) in the unpaced segment is visible.

**Figure 2:**
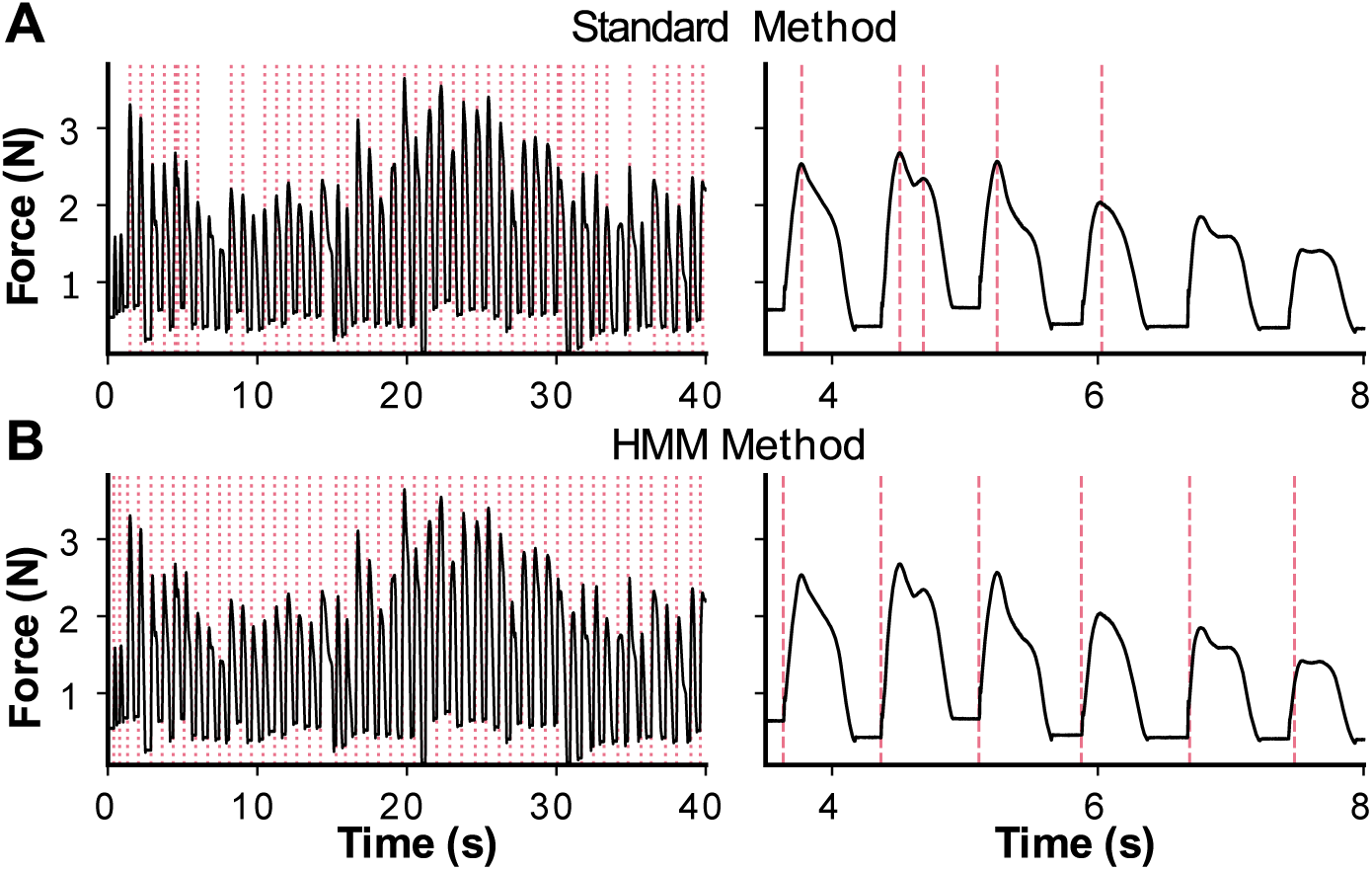
Comparison of the performances on tap detection between two analysis methods. **A:** Applying the standard method using a customary peak finder function. **B:** Using the Gaussian Hidden Markov Model (GHMM) shows how marking double peaks is avoided and how also the onsets of lighter contacts can be reliably detected.

### Spontaneous Motor Tempo (SMT)

A first goal was to measure the spontaneous motor tempo of each volunteer. A custom-written algorithm calculated the mean SMT and the 20% faster and 20% slower periods and customized the pacing protocol for the immediately ensuing data collection. Figure 3 shows the preferred periods as a function of age, separated for individuals with and without musical experience. The SMT was on average 528ms, which is close to 500ms, ubiquitously reported in the literature as the preferred rate for production^34,9,37^ and perception^9^.

**Figure 3:**
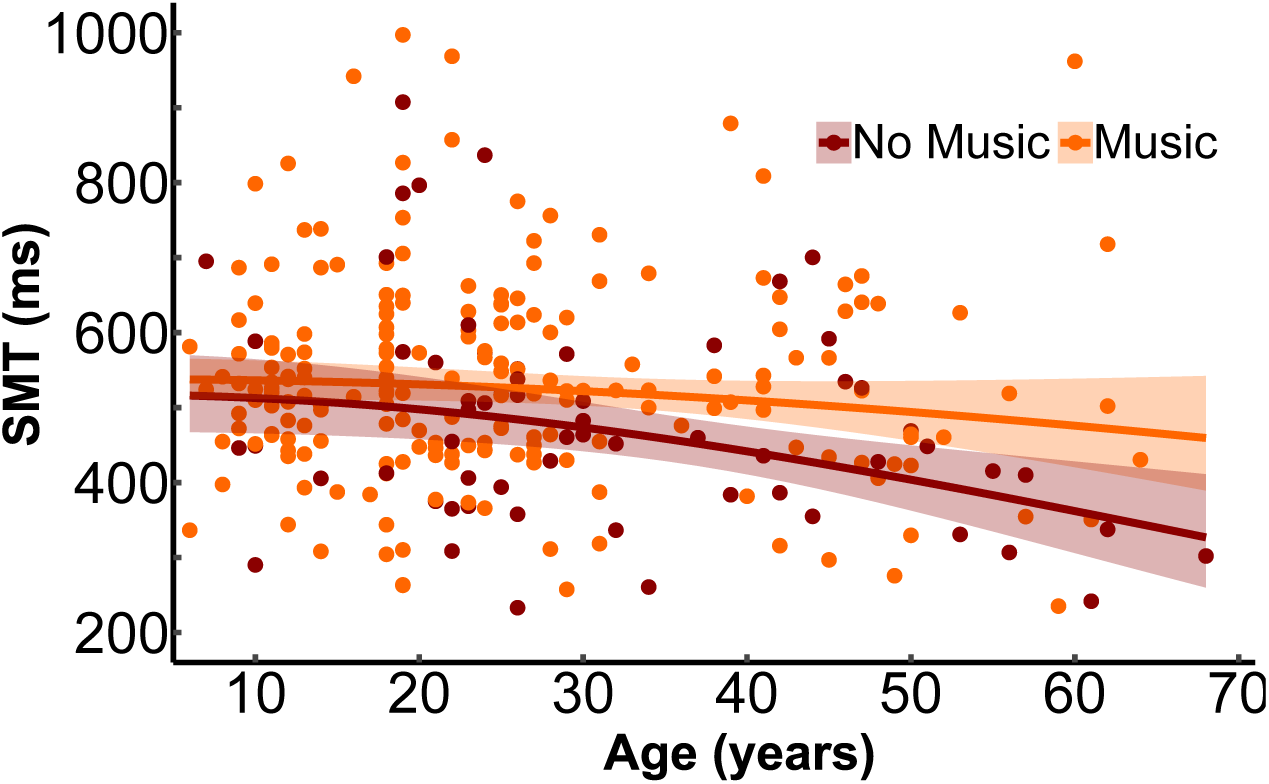
Spontaneous motor tempi (SMT) as a function of age split by musical experience. One point represents one participant (SMT average over one trial). The solid lines indicate the parabolic regression, separated by musical experience. Shaded areas represent the 95% confidence interval (estimate -/+1.96 SE).

Yet, it also showed wide variations between 233ms and 1122ms, partly due to the uncontrolled environment, partly to the wide range of individuals. Musical experience significantly affected the SMT (LRT=78.82, df=1, p=0.003), showing that it was on average 49ms longer for subjects with musical experience (577ms, t=2.41, p=0.016). The interaction between age (squared) and musical experience showed a weak tendency for significance (LRT=2.77, df=1, p=0.096): as seen in Figure 3, the SMT tended to decrease with age, but only for subjects with no musical experience (beta= -9.95e-05, t=-3.036, p=0.005).

### ITI Error

The error of the intertap (ITI) interval was given by the median of the unsigned deviation of the actual tapping period from the metronome period. It was expressed as percent of the target duration to allow pooling all different periods. The main effects of age (β=-0.023, LRT=17.51, df=1, p<0.0001), age squared (β=0.00028, LRT=12.16, df=1, p=0.0005), musical experience (LRT=10.85, df=1, p=0.001) and the interaction between segment and period condition (LRT=16.56, df=2, p=0.0003) were significant. Figure 4A shows the errors against age, with a parabolic regression, suggesting that timing error decreased from children to adults and increased again at older age. The paced segment showed overall smaller errors than the unpaced continuation segment. Figure 4B highlights how musically experienced subjects exhibited a lower timing error than subjects without musical experience (music–no music=-0.14, t=3.32, p=0.001). Figure 4C showcases how the unpaced segment had significantly higher timing error than the paced segment (paced–unpaced=-0.57, t=-24.17, p<0.0001). Within the paced segment, the faster condition exhibited higher timing errors than the preferred condition (β=0.14, t=4.63, p<0.0001) and the slower condition (β=0.14, t=4.74, p<0.0001). This is consistent with the oscillator hypothesis.

**Figure 4:**
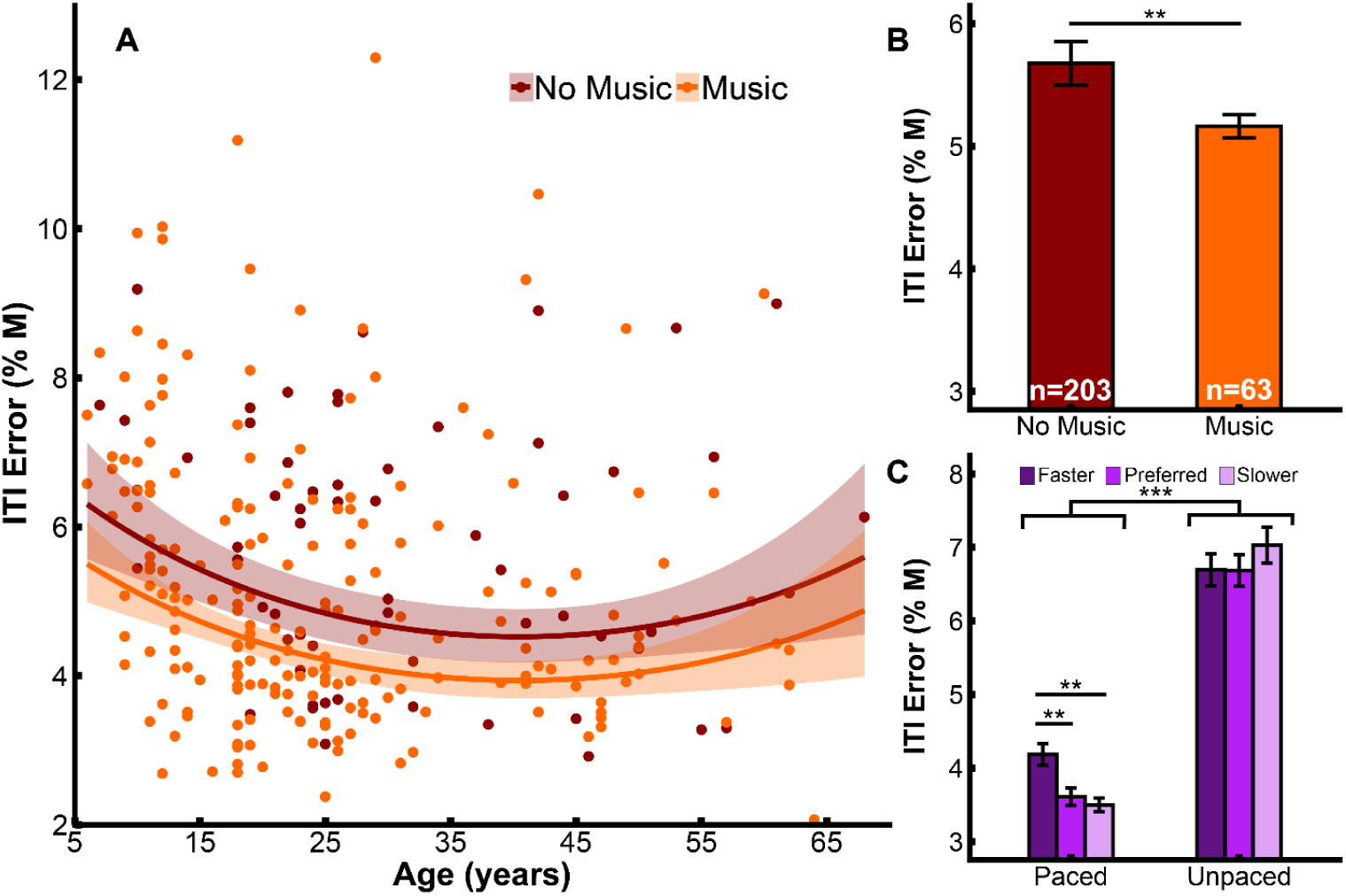
Timing (ITI) error (in percent of the metronome period M) as a function of age, music experience, segment and period condition. **A:** Timing error as a function of age. One point represents one participant averaged over all conditions and segments. The solid lines indicate the parabolic regression, separated by muscial experience. Shaded areas represent the 95% confidence interval (estimate -/+1.96 SE). **B:** ITI error for participants with and without musical experience, pooled across all period conditions. The total number of participants in each group is given in the bars. **C:** ITI error in the paced and unpaced segments separated by the three period conditions. Brackets indicate significant differences with their corresponding level of significance (* p<0.05; ** p<0.01; *** p<0.001). Error bars represent the standard error.

### Asynchrony

The asynchrony was the signed difference between the metronome and its corresponding tap onset, with negative values indicating that the tap advanced the metronome. This metric was only computed for the paced segment. As Figure 5 shows, it was indeed negative across all ages with an overall average of -19ms. It was mainly modulated by music experience (LRT=10.24, df=1, p=0.001) and period condition (LRT=98.87, df=2, p<0.0001). Figure 5A shows a tendency for asynchrony to increase with age (β =− 0.0009, LRT = 3.37, p = 0.066). Figure 5B shows how asynchrony was longer for individuals without musical experience (music-no music=3.51, t=3.23, p=0.001). As seen in Figure 5C, the asynchrony decreased as the target interval changed from the faster to the slower periods (faster–preferred=2.22, t=4.82, p<0.0001; faster–slower=4.82, t=10.35, p<0.0001; preferred-slower=2.60, t=6.75,p<0.001).

**Figure 5:**
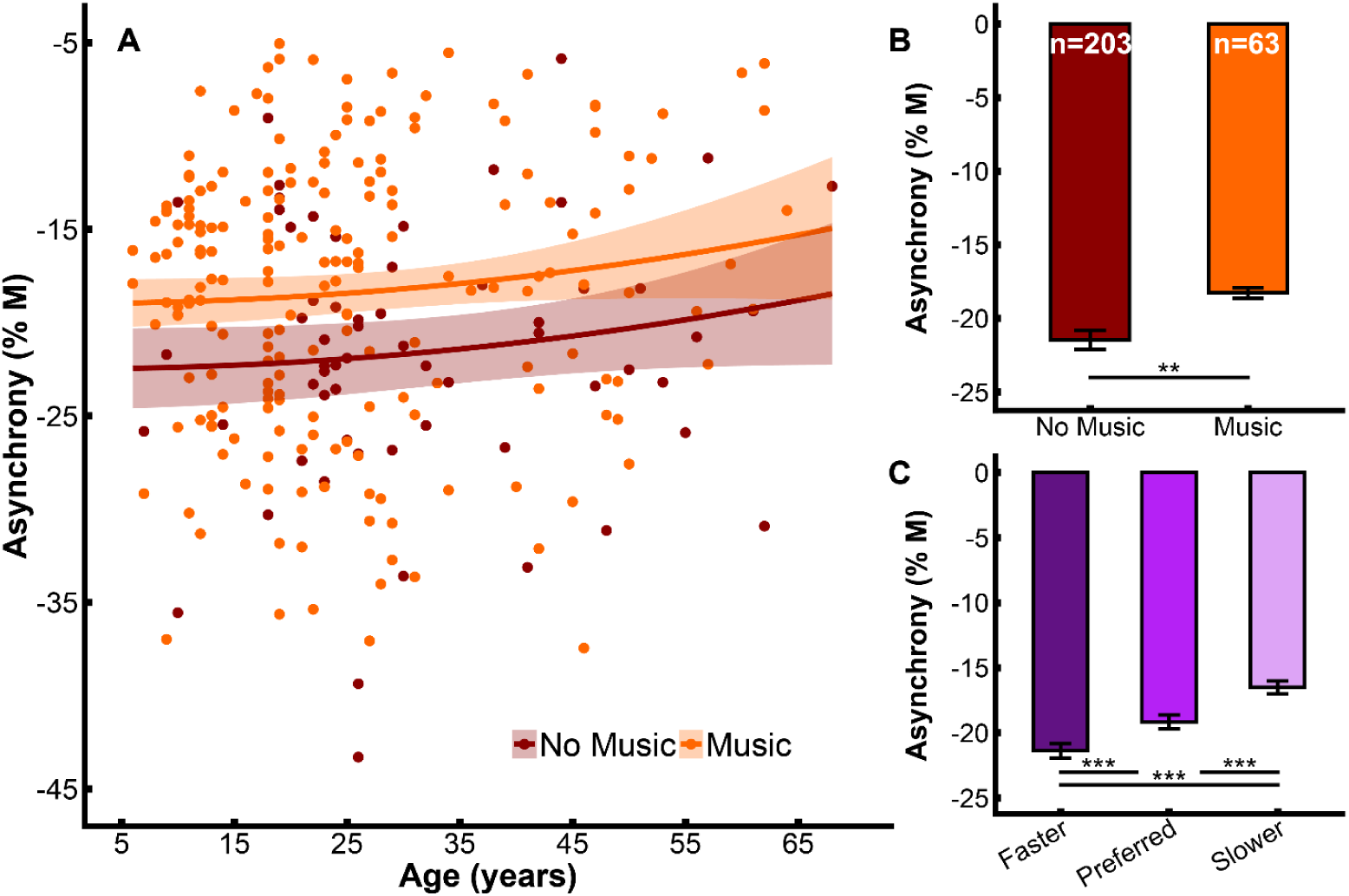
Asynchrony as a function of age, music experience, and period condition. **A:** Asynchrony as a function of age, separated by musical experience. One point represents one participant averaged over all period conditions on the paced segment. The solid lines indicate the parabolic regression, with and without music experience. Shaded areas represent the 95% confidence interval (estimate -/+1.96 SE). **B:** Asynchrony for participants with and without musical experience, pooled for all period conditions. **C:** Asynchrony in the three period conditions, pooled over music experience. Brackets indicate significant differences with their corresponding level of significance (* p<0.05; ** p<0.01; *** p<0.001). Error bars represent standard error.

### ITI Variability

The ITI variability was defined by the quartile variation coefficient. Similar to the ITI error, age (β=-0.02, LRT=29.13, df=1, p<0.0001), age squared (β=0.0003, LRT=16.81, df=1, p<0.0001), musical experience (LRT=10.57, df=1, p=0.001) and the interaction between segment and period condition (LRT=8.50, df=2, p=0.014) had significant effects on ITI variability. Figure 6A shows how ITI variability first decreased and then increased with age, expressed as a function of age-squared. Musically experienced participants displayed lower variability (music– no music=-0.11, t=-3.28, p=0.001; Figure 6B). As seen in Figure 6C, in the paced segment, ITI variability was higher in the faster condition compared to the preferred (faster–preferred=0.1, t=3.68, p=0.0007) and the slower conditions (faster-slower=0.095, t=3.57, p=0.001). In the slower and preferred conditions, the unpaced segment showed higher variability than the paced segment (slower: paced–unpaced=-0.11, t=-4.29, p<0.0001; preferred: paced–unpaced=-0.15, t=-4.77, p<0.0001). This is consistent with an oscillator interpretation.

**Figure 6:**
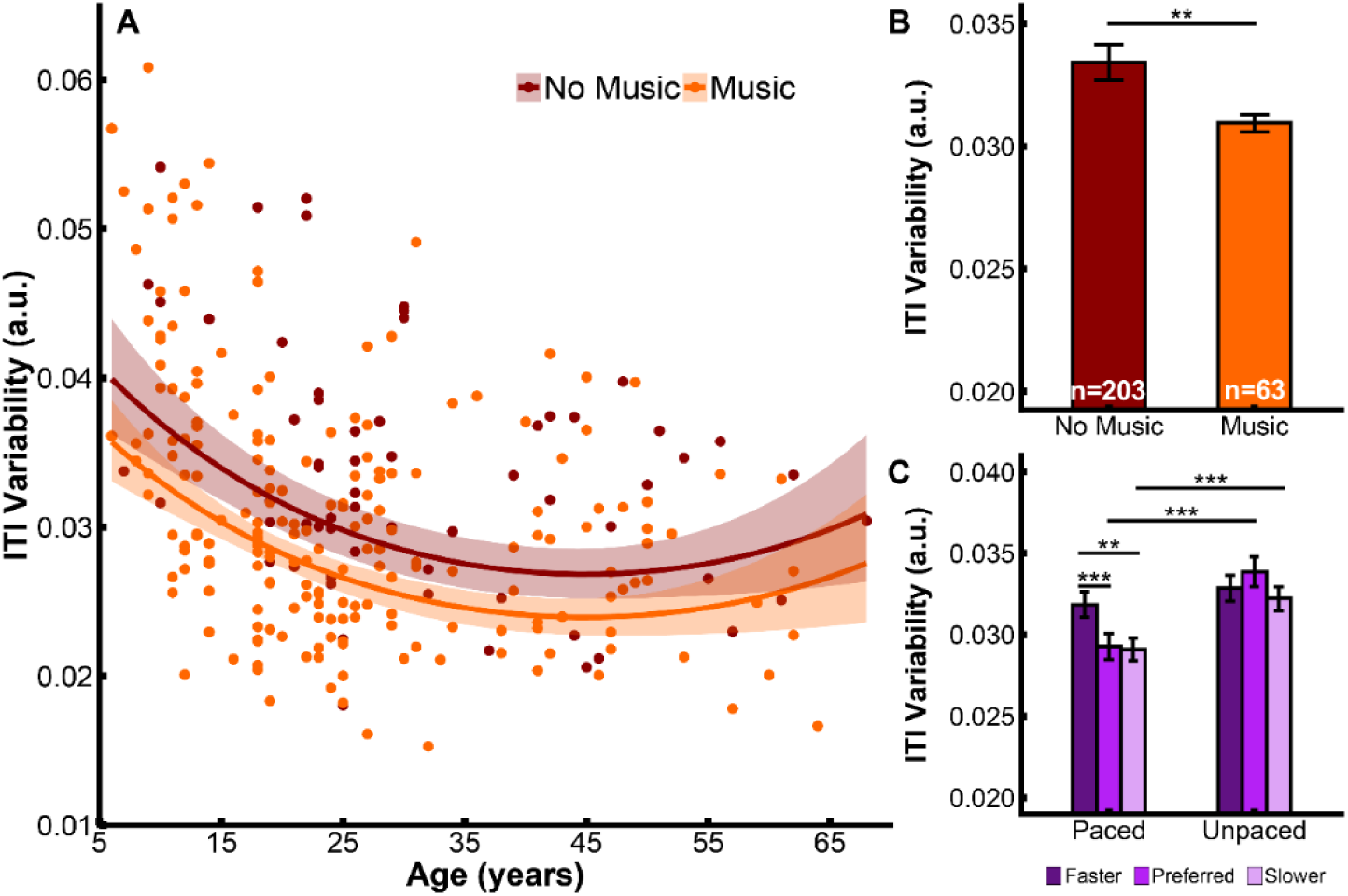
ITI variability as a function of age, musical experience, segment and period condition. **A:** ITI variability as a function of age, split by musical experience. One point represents one participant averaged over all conditions and segments. The solid lines indicate the parabolic regression, with and without muscial experience. Shaded areas represent the 95% confidence interval (estimate -/+1.96 standard error). **B:** ITI variability in participants with and without musical experience, pooled for all period conditions. **C:** ITI variability in the paced and unpaced segments separated by the three period conditions. Brackets indicate significant differences with their corresponding level of significance (* p<0.05; ** p<0.01; *** p<0.001). Error bars represent standard error.

**Figure 7:**
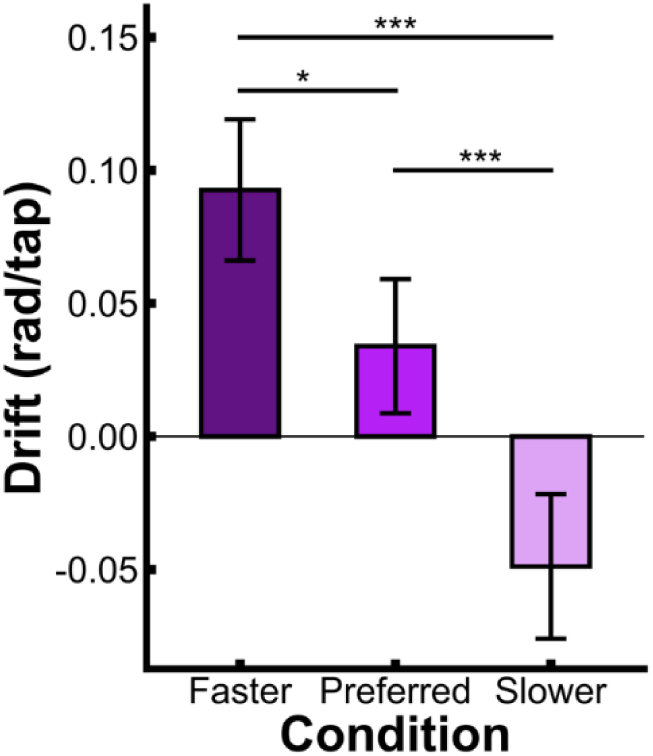
Drift as a function of period condition. Brackets indicate significant differences with their corresponding level of significance (* p<0.05; ** p<0.01; *** p<0.001). Error bars represent standard error.

### Drift

Expressed in radians per tap, the drift can be negative or positive, depending on whether the participant tapped faster or slower than the initial period. As hypothesized, drift significantly differed in the three period conditions (LRT=36.56, df=2, p<0.0001): drift was significantly positive in the 20% faster condition (β=0.11, t=4.37,p=0.0001), showing that subjects slowed down their tempo when they were no longer paced by the metronome. The drift was also significantly higher in the faster period than in the preferred and slower period (faster– preferred=0.082, t=3.91, p=0.0003; faster–slower=0.16; t=6.23, p<0.0001). As expected, in the slower condition drift was negative, reflecting the tendency to speed up, but this effect was not significant (β =-0.043, t=-1.58, p=0.3). However, this trend was reinforced by the drift being significantly lower in the slower than the preferred condition (preferred–slower=0.074; t=3.29, p=0.003). Counter to our hypotheses, neither age, nor musical experience significantly impacted the drift.

## Discussion

Over more than 100 years, legions of experiments have employed the extremely simple movement – finger tapping - to test the ability to react, predict and entrain to auditory or visual stimuli. The synchronization-continuation paradigm, first introduced by Stevens in the 19^th^ century, has accumulated numerous studies and a range of robust results. Motivated by the desire to add ecological validity to the fine-grained results on motor timing, this study tested this widely used protocol in a more naturalistic setting. From the vantage point of dynamical systems theory, we designed pacing periods anchored in the preferred period of each individual participant, which was measured directly before the data collection proper. We hypothesized that this preferred period acts as an attractor and, when moving at non-preferred rhythms, elicits not only higher error and variability, but also drift towards the preferred period after having been paced at a different period.

### From the Controlled Laboratory to the “Living Laboratory”

The body of work on rhythmic timing has accrued a set of well-established findings obtained in rigorously controlled experiments, with individuals that were motivated and frequently rewarded. The metrics that describe timing acuity are in the order of milliseconds, and the effect sizes are often much below 100ms. What do these results mean for behavior in the real world? Our first step to address this question was to repeat the experimental paradigm in an uncontrolled “living laboratory”, involving a wide variety of individuals with different ages, educational and cultural backgrounds.

Given this wide variety spectrum of participants, the pacing conditions in our study were anchored in the spontaneous or preferred motor tempo, with other periods scaled to it. This experimental design is grounded in the assumption that our brain does not appreciate arbitrary external values (except may be people with absolute pitch), but perceives and performs external tempi relative to its own intrinsic pace. This assumption comports with Gibsonian tenets that humans measure space and time in intrinsic coordinates^38^. From a dynamic systems perspective, every physical system when oscillating has a natural frequency that has the highest degree of stability and therefore requires the least amount of control or energy. Although a tapping finger has little inertia compared to, for example, a swinging arm, a natural frequency exists when moving in the gravitational field. Although several studies have investigated SMT as a reliable phenomenon, e.g.^39^, our study went one step further and investigated synchronization to external tempi in the vicinity of this intrinsically preferred period.

### Robust Results

The study took place in a museum exhibition hall and included more than 300 participants between 4 to 68 years old with diverse musical backgrounds. This musical experience included all instruments and ranged from nothing to 40 years, typically confined to leisure activities (with some exceptions). The results replicated the main findings from the literature: (1) subjects exhibited an SMT close to 500ms; (2) ITI error and variability and asynchrony decreased during childhood and increased again in older people; (3) individuals with musical experience showed less error and variability and slightly slower preferred periods.

### Preferred Periods as Natural Periods of a Dynamical System

A central result for our hypothesis was that when paced at rhythms different from their SMT, individuals drifted back to their preferred tempo when the external pacing stopped. These findings supported the hypothesis that the preferred period acts as an attractor, i.e., timing is not absolute, but individually grounded. Note that the SMT varied considerably between participants, ranging from 300ms to 1000ms. Natural periods have been recognized as inherent to locomotion, with the preferred stride frequency accounted for by biomechanical and energetic factors^40–42^. In this same dynamical interpretation, Yu and colleagues^7^ revealed a drift towards the natural period in a manual task. However, this task involved swinging different hand-held pendulums that had significant inertia and suggested biomechanical explanations. When tapping a finger, biomechanics is not as prominent as a predictor of the preferred tempo as the finger’s inertia is almost negligible. Hence, our observed drifts towards the preferred periods support that also neural considerations come into play.

In the early 2000’s, some research reported evidence in favor of the dynamic systems interpretation as an alternative to the prevailing information processing model that distinguished between a clock component and additive stochastic noise^7,9,17,43,44^. Sternad and colleagues^7,43^ showed that ITI variability followed a U-shape with its minimum at the SMT, against the idea of variability being proportional to the interval duration. Our findings also showed that moving at the preferred period exhibited the smallest drift. This result fits well with conceptualizing rhythmic timing as an oscillator at its natural frequency.

### Asymmetry in Sensorimotor Timing

Previous research showed that the range of periods that humans can tap without significant deterioration is from 150ms to 1500ms. After 1500ms, beats are no longer anticipated and asynchrony becomes positive, i.e., taps react to the beat^12^, and ITI variability increases. In our dataset, 25% of the trials in the faster period condition were between 233ms and 437ms, with a mean at 361ms. Hence, the SMT of 528ms is closer to the fastest possible tapping pace than to the limit for slow movements^12,45^. Again assuming that the SMT is a neuromechanical attractor, the basin of attraction is quite asymmetric. The hypothesis of an attractor with an asymmetric basin of attraction received some support by results within the paced segments: 1) intervals 20% faster than the SMT displayed a higher error and variability than the slower intervals; 2) the drift to the SMT was more pronounced from the faster periods than from the slower periods.

### Changes with Age

In line with previous work, ITI error and variability showed a U-shape across the age range; in addition, asynchrony decreased with age. These results were seen in all three period conditions, including the SMT, suggesting that timing at younger and older ages is subject to general developmental and aging changes. While drift was previously shown to decrease from childhood to adulthood^9^, our results did not replicate these findings. Firstly, our dataset only included 8 children under 8 years old. Then, periods were confined to 20% away from the SMT; age differences may have only shown up for periods further away. More importantly, the discrepancy may be due to different calculation: McAuley and colleagues^9^ quantified drift as the average error over all taps during continuation for target intervals from 150ms to 1700ms. Our analyses quantified the drift by relative phase in the last converging portion of the continuation segment. This choice recognized the drift as a non-stationary process, following earlier results that had shown that drift was an exponential return to the preferred period^7^, consistent with the behavior of a first-order dynamical system.

### Musical Experience

In line with previous work^28^, ITI error, variability and asynchrony were lower for participants with some musical experience, suggesting that musical training increases the flexibility to entrain to other non-preferred rhythms. However, musicians did not perform significantly better at maintaining a steady tempo without external cues, counter the expectation that attractor stability might increase in musicians. In a similar vein, Zamm et al.^5^ reported that musicians’ tapping tempo drifted to their natural frequency when the external cue disappeared, although that study did not compare musicians with non-musicians.

One important consideration in reviewing results on ‘musicians’ is that the label has been given to individuals of very different experience and background. Dalla Bella and colleagues (2024) defined a musician as having at least 7 years of formal training, while Scheurich and colleagues (2018) reported that a couple of years of systematic training could already significantly improve sensorimotor timing. However, music training of beginners has not rendered coherent results^32,46^. In our study, the number of years of practice ranged from 1 to 40 years, with a median at 7+/-8 years. 25% of our ‘musicians’ practiced for less than 3 years. Importantly, we did not require systematic or even professional training. And yet, even with this moderate music experience sensorimotor timing was improved. Note also that the experience that our museum participants reported included basically all instruments including singing. This finding is reinforced by a recent study that investigated the heterogeneity of profiles in musicians and non-musicians^47^. They found that informal music experience among non-musicians significantly enhanced motor and perception performances. Considering this multi-dimensional continuum of music practice may be more appropriate than the binary discrimination between musicians and non-musicians applied in most studies on timing.

### Limitations

While our study recruited a large variety of participants, we could not target specific individuals. Hence, the age distribution was uneven, with fewer children and elderly than middle-age adults, weakening the generalizability of the findings across the life span. Due to the same recruitment limitations, musical experience was highly variable. Nevertheless, both age and music experience had significant effects on performance, highlighting the robustness of these results. The SMT was assessed prior to the experiment using only a single trial, impacting the robustness of this metric. Due to practical constraints, we were unable to implement a more fine-grained and reliable method for assessing the SMT, such as the approach proposed by Kaya and Henry^39^. Further, our living laboratory comes with practical constraints. To keep the experiment within the permitted 15min, there were only two trials per condition, and the analyzed segments were too short to thoroughly study non-stationnary processes such as the drift. Future studies would benefit from a more detailed characterization of individual musical background, a greater number of trials, and a more robust assessment of SMT across multiple sessions.

## Conclusions

Our study design was motivated by a dynamical systems perspective with the hypothesis that rhythmic timing represents a neuromechanical oscillator that presents an attractor with a basin of attraction. Several results were consistent with this hypothesis, particularly the drift observed during the continuation segment. Our explicitly uncontrolled study was also able to replicate the main previous results on sensorimotor timing, giving them ecological validity allowing some conclusions and applications in the real world. As even short-term music experience positively affected sensorimotor timing, some music training should be fruitful to improve sensorimotor timing in children. Clinical work with Parkinson patients already showed that music-based gait training not only improved gait parameters, but also perceptual and motor timing performance^48,49^. This speaks to providing musical training for neuro-diverse populations, such as autism^50^. Considering the rapid development of portable and adaptive technologies^51^, there is an opportunity to exploit these effects^36^.

## Methods

### Participants

A total of 335 volunteers participated in the study (180 female, 155 male, ages 5-68). The inidivudals were of all educational, professional, cultural and ethnic backgrounds. The study was approved by the Institutional Revew Board of Northeastern University and the Museum of Science in Boston. All subjects gave consent prior to participating in the study and nobody received any remuneration. Everyone also completed a one-page survey that collected demographic attributes, any known physical and neurological disorders, and information regarding previous musical experience.

The experiment was part of the Living Laboratory program of the Museum of Science in Boston, an initiative designed to educate the public through active involvement in scientific experiments. All subjects participating in this experiment were visitors at the Museum of Science in Boston, Massachusetts, that were attracted to the outreach efforts by a collection of students of Northeastern University. Data collection occurred during the 4-hour exhibitions that were set up every weekend between October 2015 and May 2016. The experiment was conducted by a total of 9 experimenters, including 6 undergraduate students, 2 graduate students, and 1 postdoctoral fellow. Each data collection session required the team work of 3 students—one for recruiting museum patrons, one for providing general information about ongoing research, and one for operating the experimental apparatus for data collection. The 9 students rotated their responsibilities each weekend. To ensure that they all followed the same procedures, they all underwent a streamlined introduction. The total time of data collection for each subject was not to exceed 15 minutes per museum policy.

### Data Elimination

Given the uncontrolled nature of the experimental data collection and because the experimental apparatus had to be set up every data collection session, the subjects’ data had to be closely scrutinized and several subsets of data had to be eliminated from further analysis, ending with a total of 266 subjects (149 females, 117 males, ages 6-68): based on visual scrutiny of the signals, 11 subjects were removed due to incorrect execution of the task. these subjects either tapped too lightly on the force sensor, were not able to synchronize to the auditory metronome sometimes due to distraction, or forgot to continue tapping after the auditory metronome was silenced. An additional 20 subjects were excluded during the course of post-processing due to data acquisition and processing errors. Specifically, 5 subjects were discarded because the experimental apparatus had been improperly configured. The remaining 15 of these 20 aforementioned subjects were discarded because there was too much noise within the digital signal to properly identify tap initiations. An additional 22 subjects were excluded prior to statistical testing due to abnormalities related to their SMT. Specifically, 1 subject had a SMT of 1.8 s that differed by more than 5 standard deviations from the average SMT. The other 21 subjects were discarded because the SMT that was calculated at the start of the experiment differed by more than 10% from the value obtained later. The 10% threshold was chosen based on empirical measurements of the SMT variability in the prior literature (Collyer, Broadbent, & Church, 1994; McAuley et al., 2006). Sixteen additional subjects consistently presented outliers defined below (see section ‘Dependent measures’).

Following all these eliminations, a total of 266 subjects (149 females, 117 males, ages 6-68) were included in the final analysis. The full age and gender distribution for the analyzed subjects can be found in Figure 1A. 203 subjects declared having musical experience of at least one year. Out of these subjects, 136 subjects reported their number of years of experience being between 1 and 40 years, summarized as a median of 7 years and an IQR of 8 years. The frequency of practice per week ranged from 1 to 4 times, with a median of 3 (mean of 2.6) and an IQR of 2 (STD of 1.16). Piano playing and singing were the most practiced music activities, in 33 and 26 subjects, respectively.

### Experimental Design and Procedure

Subjects were seated comfortably at the research station within the Living Laboratory at the Museum of Science (Figure 1B). An experimenter explained the experimental task and provided a pair of noise-cancelling headphones to wear during the task. To mitigate the effect of distractions within the public data collection area, subjects were instructed to point their gaze toward the experimental apparatus. Nevertheless, siblings, parents, and friends frequently gathered around the subjects, watching but also potentially distracting the subject.

In the first stage of the experiment, subjects were instructed to tap their finger on a small force plate for 20 s at a pace that they could reasonably maintain for at least 5 min. Upon completion of this first task, the intervals between the onset of two subsequent taps (ITIs) were computed. The arithmetic mean of the ITIs during the middle 10 s served as an estimate for each subject’s spontaneous motor tempo, or SMT. Based on ths estimate, two periods were determined as 0.8 (20% faster), 1.0 (SMT), and 1.2 (20% slower) than the SMT.

In the main part of the experiment, subjects were informed that they would hear a metronome and should tap in synchrony with the metronome beat to the best of their abilities for 15s (synchronization segment). This segment was followed by another 25s when the metronome was turned off. Subjects completed a total of 6 trials, with 2 trials for each of the 3 metronome conditions. The 3 metronome periods were presented in random order. Raw data for a sample trial can be seen in Figure 1C.

### Experimental Apparatus

Tapping contacts were measured by a custom-made load cell with a strain gauge, glued on top of a beam that measured the deformation of the beam. To enhance sensitivity to the relatively low forces and deliver accurate and reliable signals, two holes were drilled into the beam under the strain gauge. The measured deformation was then converted into force at a sampling rate of 200Hz. Data was encoded using a data acquisition card with a pre-amplifier to enhance the signals (National Instruments, NI USB-6343).

To ensure that subjects tapped naturally, the experimenter did not provide explicit directions for how subjects should tap on the force plate, such that the magnitude of force, the contact duration of the tap, and the actual finger movement between taps could vary. To accommodate the maximum number of museum visitors during the assigned 4-hour data collection window, the sensor could only be calibrated only once at the beginning of each day. Consequently, standard processing algorithms for detection of the onset of the tap sometimes faced some challenges.

### Data Processing

Data was extracted from the force signals, with examples shown in Figure 2. When the signal rose from the baseline, the start of the tap was indicated. However, sometimes, the finger contact was also slightly prolonged giving rise to double-peaks confounding a clear marking of the peaks. Hence, the salient landmarks for ITI calculations were the sharp rises in the force signal. However, the continuous shape of the signal created some problems for a regular peak finding algorithm (Figure 2A).

To resolve the issue, a tap detection algorithm was developed that used a Gaussian Hidden Markov Model (GHMM). In the first step, the digital signal was passed through a high-frequency moving-average filter using segments of 50 frames binned into 50 linearly-spaced force intervals to filter out high frequencies of events. The observed signal was treated as a system with two states – a baseline state that existed while the force sensor was at rest, and an active state that was invoked by a finger tap. To infer the latent states from the observed data, the GHMM was fitted to the time series within each window, using the Viterbi algorithm (Viterbi, 1967). Under this model, a change from the baseline state to an active state represented a tap initiation. To reduce the false positive rate of tap detection, the time series was segmented into time windows at three scales – two times the specific metronome period of that trial, five times that same metronome period, and the entire trial length. Active-state inferences from each overlapping time window were required to be in agreement between these three runs. To account for variations in force amplitude within the span of a single trial, the signal within each window was scaled to be within a range of 0 to 100. Inferred tap initiations that occurred in the middle of a 5 ms span of monotonically decreasing force samples were discarded, as they were likely artifacts. Additionally, tap initiations that occurred less than 33% of the trial’s metronome period after a previously detected tap initiation were ignored.

To improve the precision of tap identification, a second latent state inference procedure using the GHMM was applied. Specifically, the digital signal was segmented into non-overlapping windows, split at the midpoints of consecutive taps that were inferred during the first pass. The signal within each window was again scaled to be within a range of 0 to 100; then a two-state HMM was used to infer the baseline and active states again. Similar to the first pass, tap initiations that occurred in the middle of a 5ms span of monotonically decreasing force samples were discarded, as were any tap initiations that occurred less than 33% of the trial period after a previously detected tap initiation. This algorithm extracted reliable tap initiations and inter-tap intervals, as illustrated in Figure 2B.

***Spontaneous Motor Tempo (SMT)*** was computed from 1 trial of 20s duration in the first stage of the experiment. The SMT was defined as the average duration (in ms) between tap interval of the 20s trial.

### Dependent Measures

Four metrics were calculated to evaluate the rhythmic timing ability of subjects: ITI error, asynchrony, ITI variability, and drift. For the continuation segment, the first 3s were excluded to eliminate transients. Since the minimum number of taps within a segment was 8, all measures were computed on the last 8 tap cycles to ensure the same number of cycles across the 3 conditions and across subjects.

***ITI Error*** was defined as the median of the absolute differences between each ITI and the trial’s metronome interval; for the unpaced segment, the metronome was continued virtually to present a reference to calculate the ITI error. As the target interval varied widely between subjects and is known to scale with the metronome period, the ITI error was normalized and expressed as percent of the metronome interval M:

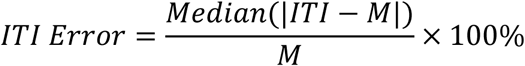

***Asynchrony*** was computed by substracting the timestamp of the metronome from the timestamp of its corresponding tap onset, and then taking the median of the differences. To discard trials where participants re-acted to the beat, i.e., tapped after the beat instead of anticipating the beat, the closest tap before a metronome beat was used as the corresponding tap. If the corresponding tap was further than half the target interval apart, this tap was considered as a syncopation, and this couple (beat and tap) was omitted from the analysis. If fewer than 4 beat-tap couples remained to compute the median, the trial was discarded. 77 trials had to be discarded. Since asynchrony is specific to the synchronization task, it was only computed for the synchronization segment.

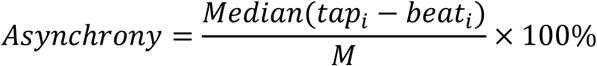

### ITI Variability

The variability of the ITIs for the synchronization and continuation segments respectively was calculated using the quartile variation coefficient *QVC_ITI_*:

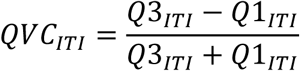

where Q1_ITI_ and Q3_ITI_ were the 25^th^ and 75^th^ percentiles of the ITI, respectively.

***Drift*** in the continuation segment was computed by first calculating the time difference between the metronome beat and the corresponding tap_i_. The time difference between the tap_i_ and its corresponding metronome beat_i_, normalized to the metronome period, was expressed as phase in radians:

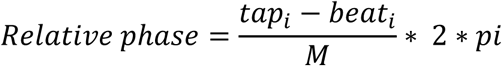

The drift was then defined as the slope, estimated by a linear regression,of the relative phase over the eight last taps

### Statistical Analyses

The study was designed to test the influence of four factors: period condition, segment, age, and musical experience. The dependent measures were ITI error, asynchrony, ITI variability, and drift. Prior to statistical analysis, outliers in the data were eliminated based on ITI error and ITI variability: trials which displayed a value larger than the 99% quantile were discarded. Based on the remainning trials, participants missing one or more conditions were deleted. This procedure removed 16 subjects from analysis (included in the counting of the eliminated subjects above). Trials were averaged for each subject in each condition and segment. Table 1 summarizes the distribution of the measured variables.

**Table 1:**
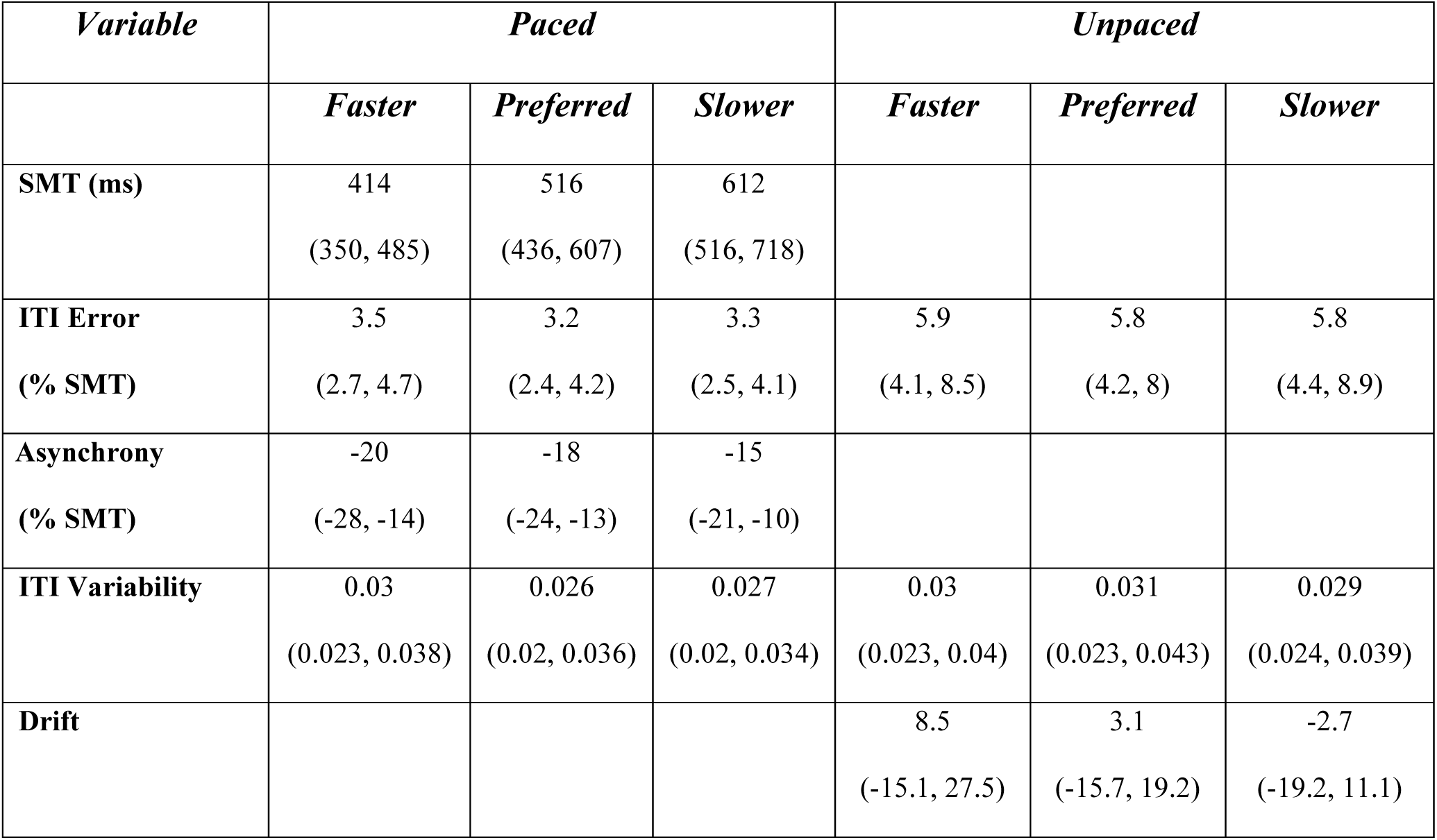
Distributions of the computed timing variables (median and first and third quartiles) for the three metronome conditions and the two segments. They are reported in milliseconds and in percentage of the SMT, with the exception of drift and ITI variability that are reported in arbitrary unit.

To test the statistical effect of period condition, segment, and musical experience as categorical variables, and age as a continuous variable, a linear mixed-effects model with subject as a random factor was used. Age was also added as a squared term to the model to account for potential opposite trends at the young and advanced age. The interaction between period condition and musical experience (and segment when applicable) were also added to the model. To correct for skewness in the distribution of the dependent variables (evaluated through visual inspection and the Shapiro-Wilk test), SMT, ITI error and ITI variability were log-transformed. Random slopes for each subject over condition and segment were included. Heterogeneity in the variance between different levels was corrected when these adjustments significantly improved the model.

The significance of the main effects and their interactions was assessed through a backward selection, using likelihood ratio tests (LRT) to compare nested models with and without each term of interest. Post-hoc comparisons with Tukey’s HSD correction at a 95% confidence level were conducted on the categorical variables and their interactions, if they remained in the model. Statistical analyses were run on *Rstudio* (v.4.3.2) using the function *lme* from the package *nlme*^52^. The post-hoc comparisons were run using the functions *emtrend* and *emmeans* from the package *emmean*s^53^.

## Acknowledgements

This study was supported by the National Science Foundation grant BCS-PAC-2043318, awarded to Dagmar Sternad.

Research Transparency Statement

## General Disclosures

Conflicts of interest: All authors declare no conflicts of interest. Funding: This study was supported by the National Science Foundation grant BCS-PAC-2043318, awarded to Dagmar Sternad. Artificial intelligence: No artificial intelligence assisted technologies were used in this research or the creation of this article. Ethics: This research received approval from a local ethics board (IRB#08-09-12).

## Study

Preregistration: No aspects of the study were preregistered. Materials: no study materials are publicly available but can be requested from the corresponding author. Data and analysis script: All data and analysis scripts are publicly available (https://osf.io/7ynb6/?view_only=0294af40e9104c3196f25b4f9d479264).

